# Role of mitotic diffusion barriers in regulating the asymmetric division of activated CD8 T cells

**DOI:** 10.1101/2021.09.10.458880

**Authors:** Hulya Emurla, Yves Barral, Annette Oxenius

## Abstract

Upon their activation, naïve CD8 T cells divide and differentiate into short-lived effector cells, relevant for exerting immune control, and long-lived memory cells, relevant for long-term immunity. The proportion of memory cells generated depends highly on the context of activation and whether the activated cell divides symmetrically or asymmetrically. However, how T cells control the extent of their asymmetry during their first division in response to contextual signals is not known. Using fluorescence loss in photo-bleaching (FLIP) experiments, we show that the metabolic and plasma membrane asymmetry of mitotic T cells depend on the regulated assembly of a lateral diffusion barrier in their endoplasmic reticulum (ER-) membrane. In asymmetrically dividing T cells, the degrees of asymmetry correlated tightly to barrier strength, whereas symmetrically dividing T cells did not establish such a barrier. Direct positive or negative interference with barrier assembly enhanced or abrogated metabolic and plasma membrane asymmetry, respectively, indicating that barrier strength is a direct and decisive determinant of mitotic asymmetry. Thus, together our data identify diffusion barrier-mediated compartmentalization as a mechanism for how asymmetric T cell regulate their long-term response as a function of the activatory context.

## Introduction

Antigen-induced activation of T cells entails the integration of three signals: 1. cognate antigen (peptide/MHC class I complexes) recognition by their T cell receptor (TCR); 2. co-stimulation via receptor-ligand interactions such as CD28 on T cells and CD80/CD86 on antigen presenting cells (APCs); and 3. pro-inflammatory cytokines, such as type I interferons or interleukin (IL)-12 ^1, 2^. This activation leads to clonal expansion and differentiation. The clonally expanded cells exhibit substantial heterogeneity with respect to phenotype, function, longevity, transcriptional, epigenetic and metabolic profiles (reviewed in ^3^). Not including the entire heterogeneity observed at the single cell level, these cells are often categorized as short-lived effector and memory precursor cells which give rise to long-lived memory cell, the former being endowed with potent effector functions such as cytotoxicity and pro-inflammatory cytokine expression to combat an infection, and the latter providing long-term protection in case of reinfections (reviewed in ^3^) Phenotypically, short-lived effector cells are defined by low expression of IL-7Rα and high expression of KLRG1 (KLRG1^hi^ IL-7Rα^lo^) ^3, 4^ whereas memory precursor cell exhibit high expression of IL-7Rα and are negative for KLRG1 (KLRG1^lo^ IL-7Rα^hi^) ^5, 6^. Memory CD8 T cells are maintained in an antigen-independent, IL-7 and IL-15-dependent manner, supporting their survival and self-renewal ^7^. The population of memory cells also exhibits heterogeneity, embracing subsets that vary in phenotype, function and localization ^8, 9, 10^. Conventionally, memory cells can be divided in central memory T cells (TCM, characterized as CD62L^hi^ CCR7^hi^) and effector memory T cells (TEM, characterized as CD62L^lo^ CCR7^lo^). TCM cells reside in secondary lymphoid organs and have greater proliferative potential than TEM cells, while TEM cells are found in peripheral organs and are equipped to readily perform effector functions ^9, 10^. More recently, an additional subset was identified as tissue-resident memory cells (TRM), which are permanently lodged within peripheral tissues where they can immediately respond to invading pathogens ^11^.

The diversity of cell fates that is adopted by progeny of activated CD8 T cells is not restricted to the ensemble of antigen-specific CD8 T cells but is also apparent for the progeny of a single CD8 T cell, as demonstrated by single cell adoptive transfer experiments or transfer of barcoded naïve T cells, followed by *in vivo* priming. The generation of diverse progeny from a single naïve T cell documents that fate is not pre-determined and is initiated after T cell priming ^12, 13, 14, 15^. However, how divergence of fates is achieved is still incompletely understood. A number of processes were shown to be involved, including the overall potency of activation (i.e. the combined strengths of signals 1-3), leading to differential transcriptional activity ^3, 16^, different epigenetic control of gene expression ^17, 18^, preferential use of different metabolic pathways ^19, 20, 21^, and asymmetric cell division (ACD) ^22, 23, 24, 25, 26^.

In a broad range of situations in eukaryotes, the generation of cellular diversity is supported by ACD, allowing unequal segregation of cell fate determinants between sister cells during mitosis, For instance, asymmetric division of budding yeast, *C. elegans* oocytes, Drosophila neuroblasts and mouse neural stem cells helps to generate both self-renewing and differentiating progeny through the asymmetric segregation of differentiation and ageing factors ^27, 28, 29, 30^. Likewise, activated CD8 T cells can divide asymmetrically, and this is thought to contribute to the generation of diverse progeny as the immune systems reacts against a specific threat ^22^. Particularly, asymmetric cell division favours the emergence of progenitors that are thought to support the generation of short-lived effector cells and long-lived memory precursor cells ^13, 16^. Strikingly, however, not all naïve T cells divide asymmetrically upon activation and when they do, the asymmetry of partitioned factors can largely vary between divisions. For example, in asymmetrically dividing CD8 T cells the plasma membrane protein CD8 may accumulate from 1.5 to more than 10 times more in one daughter over the other ^31^. The extent of polarity and asymmetry depends strongly on the strength and context of activation ^32^. Thus, the asymmetry of the cell during division is thought to dictate how heterogeneous the progeny of a given naïve T cell will be upon its activation and to impact the relative composition effector and memory cells ^31^. Thus, increasing evidence indicates that asymmetric cell division and its gradation helps to adapt CD8 T cell responses according to strength and context of activation. However, how CD8 T cells control whether and to which extent they divide symmetrically or asymmetrically is largely unknown.

Activation of naïve CD8 T cells by signals 1-3 induces polarization of engaged cell surface molecules (e.g. T cell receptor -TCR, CD3, CD8 and LFA-1) towards the immunological synapse that establishes between the antigen-presenting cells (APC) and the engaging CD8 T cell ^24^. Depending on signal composition and strength, this asymmetry may be maintained in mitosis, promoting the unequal partitioning of these receptors during mitosis. Additional cellular constituents that may also become unequally distributed between daughter cells include surface receptors not directly involved in the immunological synapse (IFNγR, and IL-2Rα ^24^), master transcription factors (T-bet; ^33^), intracellular degradation machineries (the proteasome; *14*), metabolic signalling hubs (mTORC1 and mitochondria; ^34^) and conserved polarity factors (scribble, Numb and PKCζ; ^24^). Accordingly, CD8 T cells that were exposed to strong activation signals (e.g. high-affinity TCR ligands, high levels of costimulation ^35^) were more likely to undergo asymmetric cell division and to generate daughter cells that diverged in their transcriptional and metabolic profile ^19, 34, 36, 37^. Together, these observations suggest that a combination of signals regulates the mitotic polarity of activated CD8 T cells to determine the asymmetry extent of their division.

Studies of the mechanisms of asymmetric cell division in yeast, tissue culture cells and in mouse neural stem cells have revealed the importance of the *endoplasmic reticulum* (ER) in the asymmetric partition of a broad variety of cellular components, ranging from protein aggregates, ubiquitinated proteins and non-chromosomal DNA, to transcription factors and other fate determinants ^38, 39, 40, 41, 42, 43, 44^. In these cells, the ER-membrane is compartmentalized in two domains through mitosis, one in each future daughter cell, by a barrier to lateral diffusion in the future plane of cleavage. These separate domains serve as platforms for the partition of aging factors and fate determinants. Such barriers have been reported for budding yeast cells, the one cell embryo of the nematode *C. elegans* and neural stem cells. At least in yeast, the formation of this barrier depends on the enrichment for long ceramides in the ER-membrane at the barrier site ^39^. This leads to the thickening of the lipid bilayer locally and prevents passage of most ER membrane proteins, the transmembrane domains of which are adapted to the thinner bilayer that forms the bulk of the ER-membrane ^38^.

Whether asymmetrically dividing T-cells compartmentalize their ER or not and whether this plays any role in their asymmetric partition of cellular factors was not known. We therefore addressed whether murine CD8 T cells may generate a lateral diffusion barrier in the endoplasmic reticulum (ER) membrane during T cell division to potentially contribute to asymmetry in partitioning of cellular cargo such as cell surface molecules or mitochondria as metabolic hubs.

## Results

### Naïve CD8 T cells can establish an ER membrane diffusion barrier during first mitosis after TCR activation

To measure the potential existence of an ER diffusion barrier in TCR activated CD8 T cells, we leveraged fluorescence loss in photo-bleaching (FLIP) to monitor the diffusion of fluorescently labelled ER membrane-integral proteins as well as ER-luminal proteins (as negative control) between future daughter CD8 T cells through first mitosis ^45^. Using lentiviral transduction of naïve CD8 T cells enabled us to express fluorescently labelled ER-membrane and ER-luminal green fluorescent protein (GFP) reporters. Specifically, we transduced naïve CD8 T cells to express a Sec61α-GFP fusion protein (MemER-GFP) as an ER membrane reporter and ER-retrieval amino acid sequence Lys-Asp-Glu-Leu (KDEL)-modified GFP (LumER-GFP) as ER luminal reporter to monitor the exchange of the ER membrane-associated and ER-luminal proteins between future daughter CD8 T cells during TCR-triggered activation and first mitosis ^45^.

First, we verified that the applied lentiviral transduction protocol enabled the expression of the GFP fusion proteins in naïve CD8 T cells and maintained a naïve phenotype in CD8 T cells before subjecting them to TCR-induced activation and FLIP microscopy. Our data confirmed that culturing naïve CD8 T cells in IL-7 and IL-15 supplemented medium allowed efficient transduction, while maintaining a naïve phenotype (Figure 1 A, B, E).

**Figure 1.**
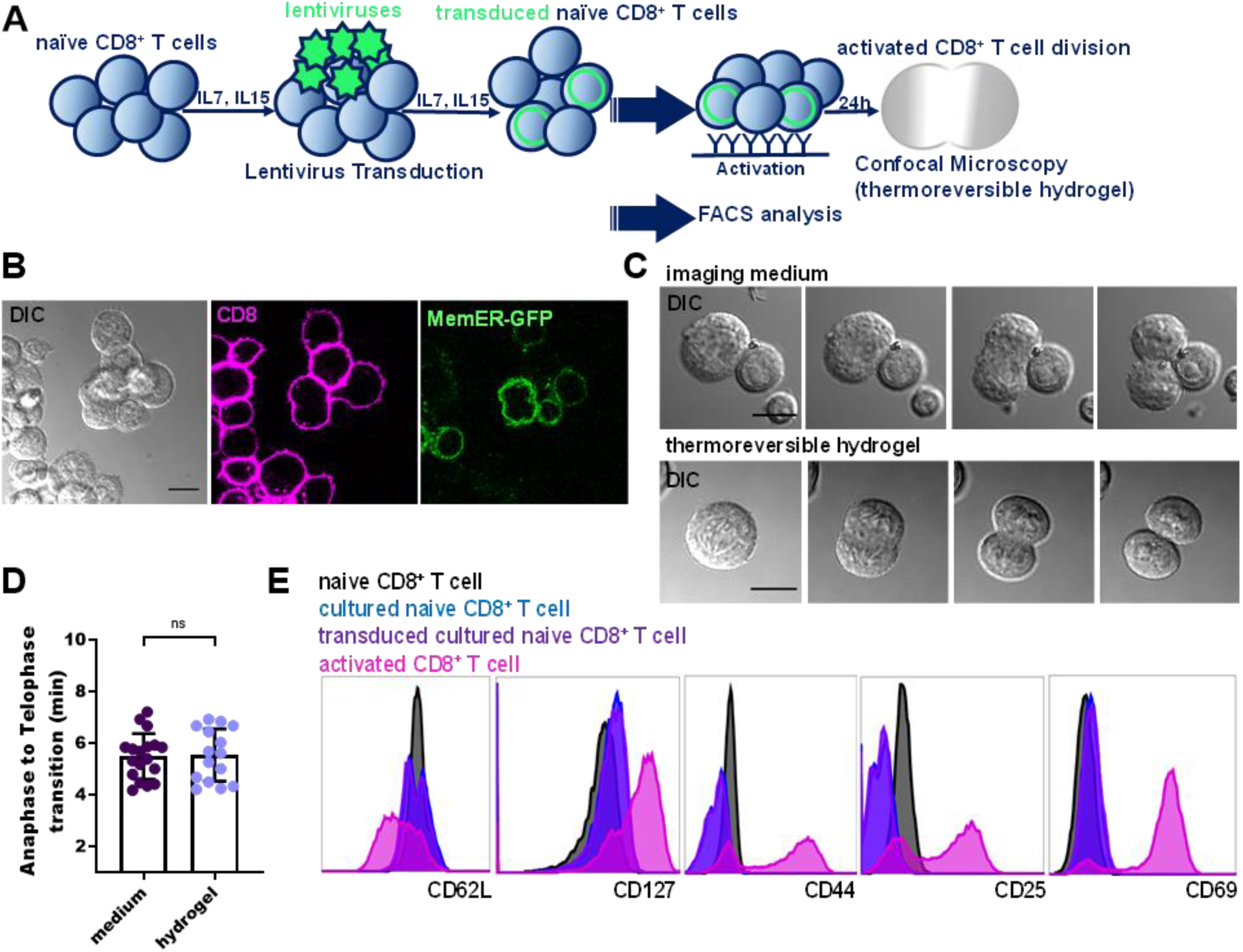
Kinetics of mitosis and phenotype of transduced naïve CD8 T cells. ***(A)** Experimental outline. Murine naïve CD8 T cells, sorted from adult mouse spleen and lymph nodes, were cultured under homeostatic proliferation condition (IL7 and IL15) and subjected to lentiviral transduction on RetroNectin coated cell culture plates, followed by activation or phenotypic analysis. **(B)** Representative images of CD8 T cells expressing* Sec61α-GFP (MemER-GFP) that stained with for CD8. ***(C and D)** Anaphase to telophase transition time of activated CD8 T cells in the imaging medium compared to thermo-reversible hydrogel. Differential interference contrast (DIC) microscopy of dividing CD8 T cells **(C)** and quantification of anaphase to telophase transitions of dividing CD8 T cells in hydrogel (blue, n=15) or imaging medium (purple, n=18) (unpaired Welch’s t-test; mean ± SEM) **(D)**. **(E)** FACS analysis of phenotypic markers (CD62L, CD127, CD44, CD25 and CD69) of lentivirally transduced (purple) or non-transduced CD8 T cells (black) prior to activation (blue), or 48 hrs after TCR-mediated activation (pink). Data are representative of three independent experiments. Unpaired Welch’s t-test; mean ± SEM. Scale bars, 10 μm.*

Second, as we embedded activated CD8 T cells in thermo-reversible hydrogel for microscopical analyses (in order to mitigate movement during mitosis), we analysed whether this would have an impact on the kinetics of mitosis (time between anaphase and telophase). We observed no difference in the time period required to transition from anaphase to telophase (Figure 1 C, D).

After having established that lentiviral transduction with the FLIP reporters would not affect the naive phenotype of CD8 T cells and that the imbedding in the thermo-reversible hydrogel would not alter the kinetics of mitosis, we identified individual early stages of T cell mitoses using differential interference contrast (DIC) microscopy in thermo-reversible hydrogel before initiating the FLIP experiments ^39^. Reporter-transduced naive CD8 T cells were activated by concomitant exposure to plate-bound agonistic antibodies for CD3, CD28 and recombinant ICAM (previously shown to promote asymmetric CD8 T cell division ^24^. Specifically, FLIP experiments were initiated at anaphase initiation until completion of telophase of mitosis. A relatively small region of interest (ROI) was repeatedly photo-bleached with a green laser through the course of T cell division, enabling to monitor comparative GFP fluorescence intensity decay in future daughter CD8 T cells (Figure 2 A). In CD8 T cells expressing the LumER-GFP reporter protein, fluorescence intensity decayed with similar kinetics in both bleached and unbleached future daughter CD8 T cells (Figure 2 D), indicating that the protein exchanged freely and rapidly throughout the cell, including across the future cleavage plane. Hence, the ER lumen is continuous in dividing CD8 T cells.

**Fig. 2.**
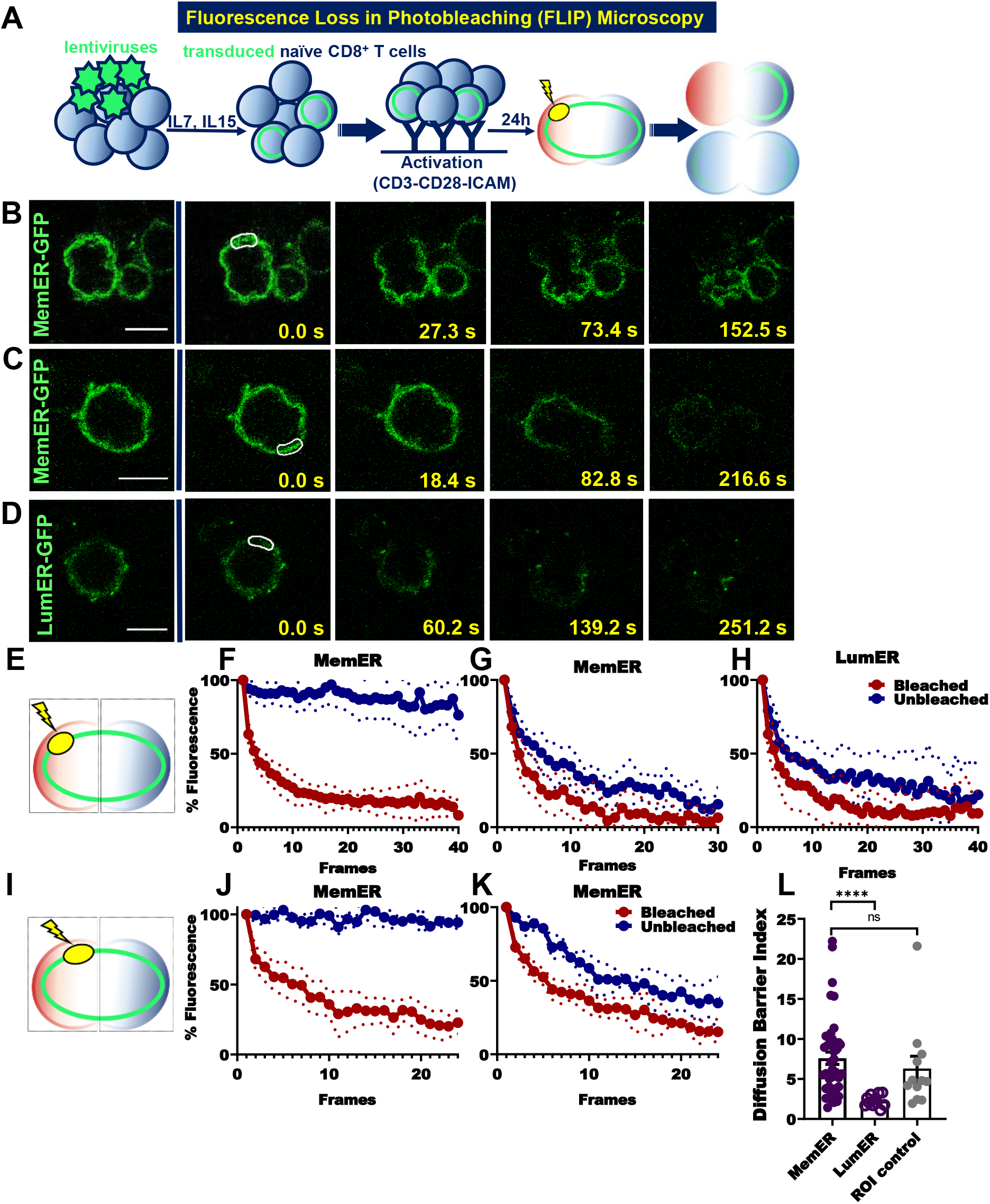
Murine naïve CD8 T cells generate a spectrum of lateral diffusion barriers in their ER membrane through their first cell division. **(A)** FLIP experiments with murine CD8 T cells that had been transduced with lentiviruses for expression of MemER-GFP **(B and C)** or LumER-GFP **(D)**. Region of interest (ROI) of repetitive photobleaching through FLIP time-lapse microscopy is indicated as a white outline. FLIP experiments of MemER-GFP results in differential compartmentalization of fluorescence intensity in the bleached future daughter CD8 T cell relative to the unbleached one, showing one example of strong compartmentalization **(B)** and one example of weak compartmentalization **(C)**. Comparable fluorescence loss in future daughter cells for LumER-GFP in FLIP experiments **(D)**. Different ROI placement relative to division plane **(E, I)**. Percentage of fluorescence intensity decay measured for both ER markers in future daughter cells with different ROI positioning throughout T cell division in bleached (red), and unbleached (blue) daughter cells for MemER-GFP (n=42), and for LumER-GFP (n=11) **(F-H, J, K).** Strength of diffusion barrier (BI: Barrier Index) quantified by nonlinear fitted curves of fluorescence decay **(L)** for MemER-GFP (filled purple circles, and grey circles), and for LumER-GFP (open purple circles); Unpaired Welch’s t-test; mean ± SEM. ****P < 0.0001. Scale bars, 10 μm.

In contrast, in a substantial (83%) fraction of CD8 T cells expressing the MemER-GFP reporter protein, the fluorescence decayed markedly slower in the unbleached than the bleached future daughter cell (Figure 2 B, E, F), indicating that in some, but not all of these cells (Figure 2 C, G), a lateral diffusion barrier compartmentalized the ER-membrane in the future cleavage plane.

In our initial experiments we positioned the ROI at a distal site of the equatorial plane of division (Figure 2 E). To verify that the positioning of the ROI does not influence the results, we repeated the experiments by placing the ROI proximal to the future cleavage plane (Figure 2 I). Positioning the bleached ROI either distal or proximal to the future cleavage plane in one of the future daughter cells did not change the results (Figure 2 E, F, I, J). Thus, these data indicate that a fraction of dividing CD8 T cells limited the exchange of the ER-membrane marker between the future daughter cells. Since the lumen of the ER was continuous throughout the dividing cell, this indicates that some, but not all, CD8 T cells assembled a lateral diffusion barrier in the ER-membrane at the future site of cleavage.

### Compartmentalization of the ER membrane during mitosis relies on TCR stimulation and PKCζ-driven polarization

To investigate whether the formation of the ER-membrane diffusion barrier is affected by specific perturbations, we next addressed whether its presence and strength is increased or decreased under specific conditions that modulate polarity or induce CD8 T cell division without involving TCR engagement (Figure 3 A).

**Figure 3.**
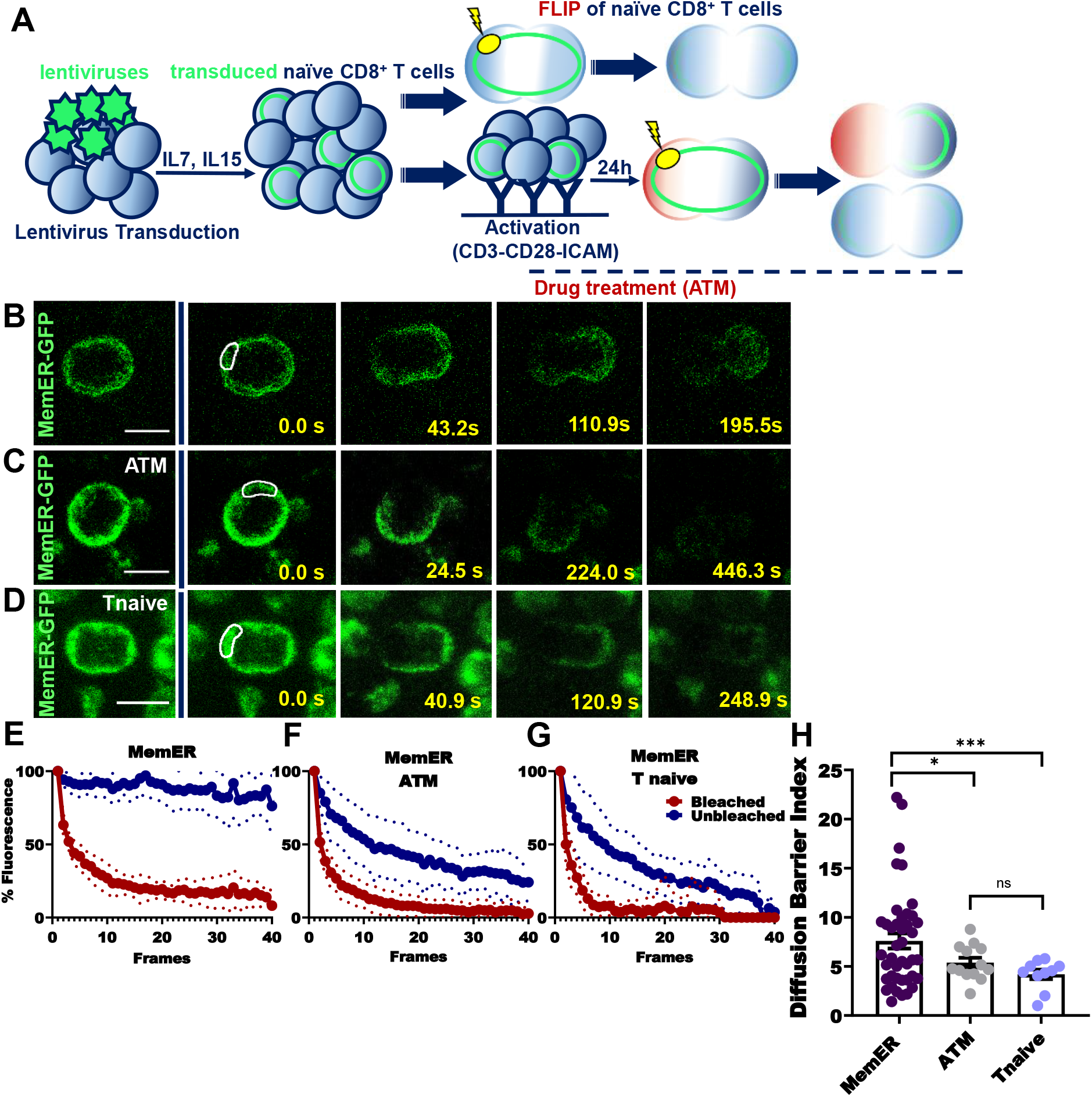
Establishment of an ER membrane diffusion barrier during mitosis of CD8 T cells relies on TCR stimulation and PKCζ-driven polarization. **(A)** FLIP experiments with murine CD8 T cells that had been transduced with lentiviruses for expression of the MemER-GFP **(B to D)**. Region of interest (ROI) of repetitive photobleaching through FLIP time-lapse microscopy is indicated as a white outline. FLIP experiments of MemER-GFP results in differential compartmentalization of fluorescence intensity in the bleached future daughter CD8 T cell relative to the unbleached one **(B)**. FLIP experiments of MemER-GFP results in comparable compartmentalization of fluorescence intensity in the bleached future daughter CD8 T cell relative to the unbleached one with polarization blockade by ATM (aurothiomalate) treatment and in naïve CD8 T cells undergoing TCR independent proliferation under homeostatic proliferation conditions **(C and D)**. Percentage of fluorescence intensity decay measured and plotted for the ER marker in future daughter cells throughout T cell division **(E to G)**. Strength of diffusion barrier (BI: Barrier Index) quantified with respect to nonlinear fitted curves of fluorescence decay (E to G) for activated T cells (purple, n=42), activated T cells treated with ATM (grey, n=13), and naïve T cells under HPC (light blue, n=10) **(H).** Unpaired Welch’s t-test; mean ± SEM. ***P = 0.0003, *P = 0.0194. Scale bars, 10 μm.

Treatment of CD8 T cells with aurothiomalate (ATM) inhibits the atypical protein kinase PKCζ, which functions downstream of the TCR to promote polarization of CD8 T cells upon their activation ^24, 46^. Strikingly, ATM treatment completely prevented barrier assembly and ER compartmentalization of CD8 T cells activated by agonistic CD3 and CD28 antibodies in presence of ICAM (Figure 3 C, F, H), indicating that PKCζ is involved in both the establishment of polarity and the formation of a diffusion barrier in the ER membrane.

Next, we asked whether induction of mitoses in CD8 T cells independently of TCR triggering would also promote diffusion barrier formation. To this end, we cultured transduced naïve CD8 T cells in conditions that favour TCR-independent cell division. Specifically, we cultured naïve CD8 T cells to in the presence of IL-7, IL-15 and IL-2 and in absence of agonistic TCR engaging antibodies. This condition induced naïve CD8 T cell divisions (homeostatic proliferation) which were then assayed for presence or absence of the diffusion barrier. Performing FLIP experiments on the membrane and luminal ER reporters of naïve CD8 T cells undergoing homeostatic cell division showed that these cells did not establish any measurable diffusion barrier (Figure 3 D, G, H). Therefore, we concluded that in CD8 T cells the compartmentalization of the ER between the future daughter cells by a lateral diffusion barrier is strictly restricted to conditions that promote asymmetric cells division, such as activation of the TCR and downstream signalling, and is influenced by the activity of polarization-promoting factors such as PKCζ.

### ER diffusion barrier strength correlates with the extent of asymmetry of dividing CD8 T cells

Next, we investigated whether barrier assembly and the mitotic asymmetry of CD8 T cells upon TCR-mediated activation were in any way correlated with each other. To this end, in addition to monitoring ER compartmentalization, we quantified CD8 asymmetry in CD8 T cells by antibody staining of CD8 post-mitotically. CD8 asymmetry has been repeatedly used for the characterization of ACD in CD8 T cells ^24, 31^. We noticed an accumulation of the CD8 signal at the contact region of sister CD8 T cells after cytokinesis, which does not allow unambiguous assignment of the signal to one or the other sister cell. We therefore decided to exclude the CD8 signal at the contact site between the two sister cells. To corroborate that this would not have an impact on the quantification of CD8 asymmetry, we compared CD8 asymmetry indices in dividing CD8 T cells with and without exclusion of the CD8 signal at the contact site between sister cells and did not find a difference (Figure S1 A, B).

We then correlated CD8 asymmetry established during TCR-triggered first mitosis with the strength of the ER-membrane diffusion barrier. Quantification of barrier strength by FLIP microscopy and polarity state by staining CD8 post-mitotically on the same cells indicated that both features correlated tightly with each other (Figure 4 A-D). CD8 asymmetry was highest for the cells that established a strong barrier (Figure 4 B, D) and lowest in those that failed to form a barrier, i.e., showed comparable loss of MemER-GFP fluorescence intensity in both daughter compartments upon FLIP (Figure 4 C, D). Quantification of barrier strength and the asymmetry of CD8 distribution demonstrated that these parameters tightly correlated with each other (Pearson correlation analysis, r: 0.9, p<0.0001) (Figure 4 D). This tight correlation suggests that assembly of a strong diffusion barrier is indispensable for activated CD8 T cells to maintain polarity during mitosis and hence, to divide asymmetrically.

**Fig. 4.**
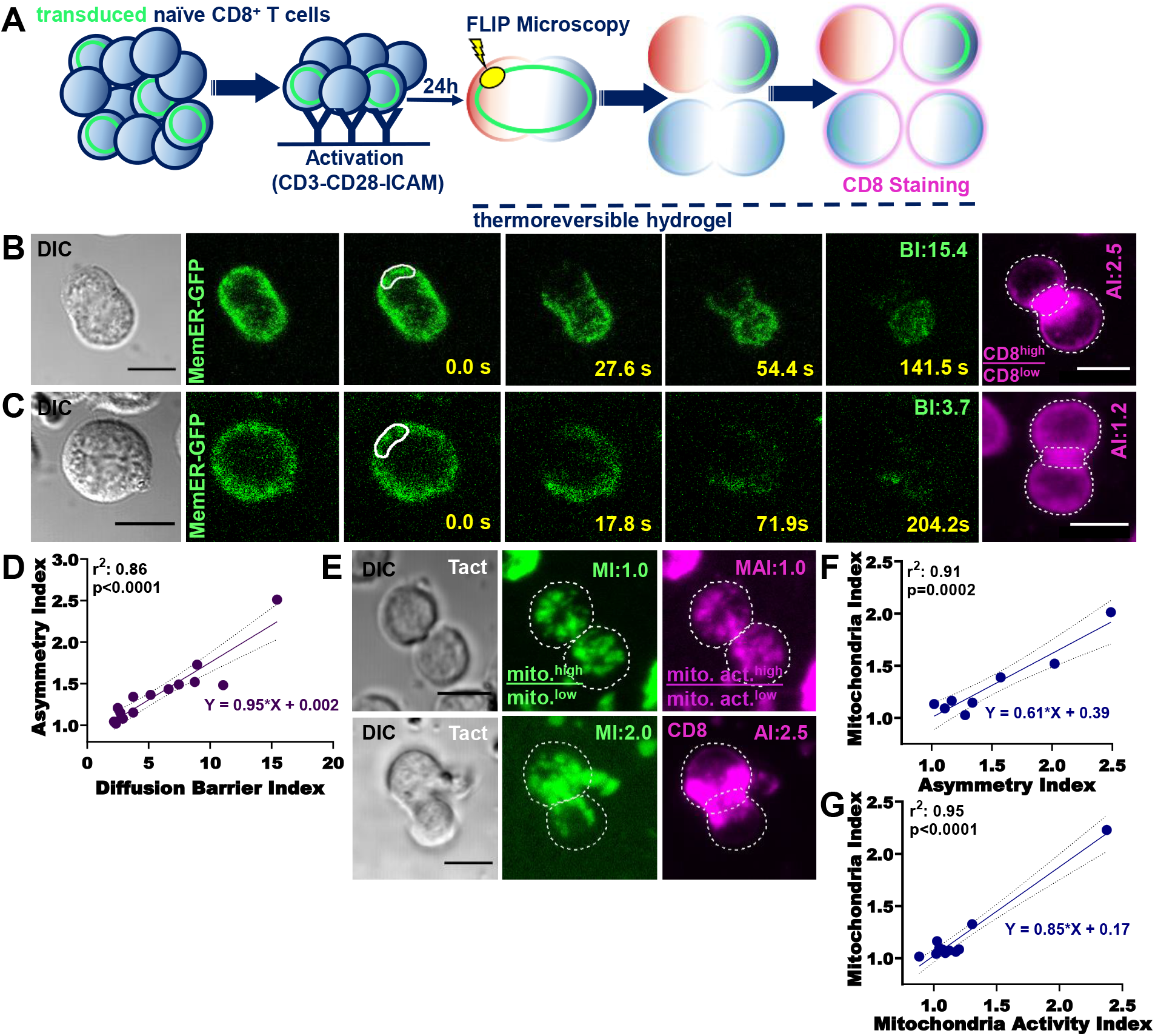
ER diffusion barrier strength correlates with the extent of asymmetry of dividing CD8 T cells. **(A)** Naïve CD8 T cells, transduced with ER reporters, were activated and subjected to FLIPmicroscopy during their first mitosis and after completion of the mitosis were stained for CD8 surface expression to delineate symmetric and asymmetric CD8 T cell divisions. Live fluorescence confocal microscopy was used to quantify the total CD8 receptor signal they inherited after completion of mitosis. Regions of interest (ROI) for repetitive photobleaching are indicated by a white outline, CD8 signal quantification was performed excluding the area that was adjacent to the cleavage plane of the dividing CD8 T cells (dashed white line). **(B)** Representative FLIP snapshots of a naïve CD8 T cell`s first mitosis that is compartmentalized via the establishment of an ER membrane diffusion barrier, giving rise to asymmetric T cell progeny. **(C)** Representative FLIP snapshots of a naïve CD8 T cell`s first mitosis that is not compartmentalized via the establishment of an ER membrane diffusion barrier, giving rise to symmetric T cell progeny. **(D)** Linear regression of the barrier index (diffusion barrier strength) and associated asymmetry index (the ratio of total CD8 signal of the sister T cell pairs) of a particular mitosis’ progeny (n=16). **(E)** Representative images of live microscopy of sister T cell pairs co-stained with MitoTracker Green (mitochondrial index: MI) and CD8 Ab, or co-stained with MitoTracker Green and MitoTracker Red (mitochondrial activity: MAI). **(F)** Linear regression of MI and MAI (n=8) from co-staining experiments. **(G)** Linear regression of MI and MAI (n=11) of co-staining (MitoTracker Red, and Green) experiments. Scale bars, 10 μm.

In addition, CD8 asymmetry also correlates tightly with the asymmetry of mitochondrial inheritance ^34^. Mitochondria were shown to be asymmetrically segregated in various other biological model systems of asymmetric cell division, such as in *Saccharomyces cerevisiae* budding ^47^, mammalian oocyte meiosis ^48^ and ACD of human mammary stem-like cells ^49^.

Quantification of mitochondrial abundance in first TCR-triggered CD8 T cell mitosis (quantified by Mito Tracker Green dye) and activity (membrane potential, measured using Mito Tracker Red) showed that both parameters correlated tightly with each other and with CD8 abundance (Figure 4 E-G). Thus, increased mitochondrial abundance correlated with CD8 abundance. In contrast, naïve CD8 T cells undergoing homeostatic proliferation in response to IL-7, IL-15 and IL-2 in absence of TCR engagement did not establish CD8 asymmetry (Figure 5B, C) and did not asymmetrically segregate their mitochondria (Figure S2 A-C).

**Fig. 5.**
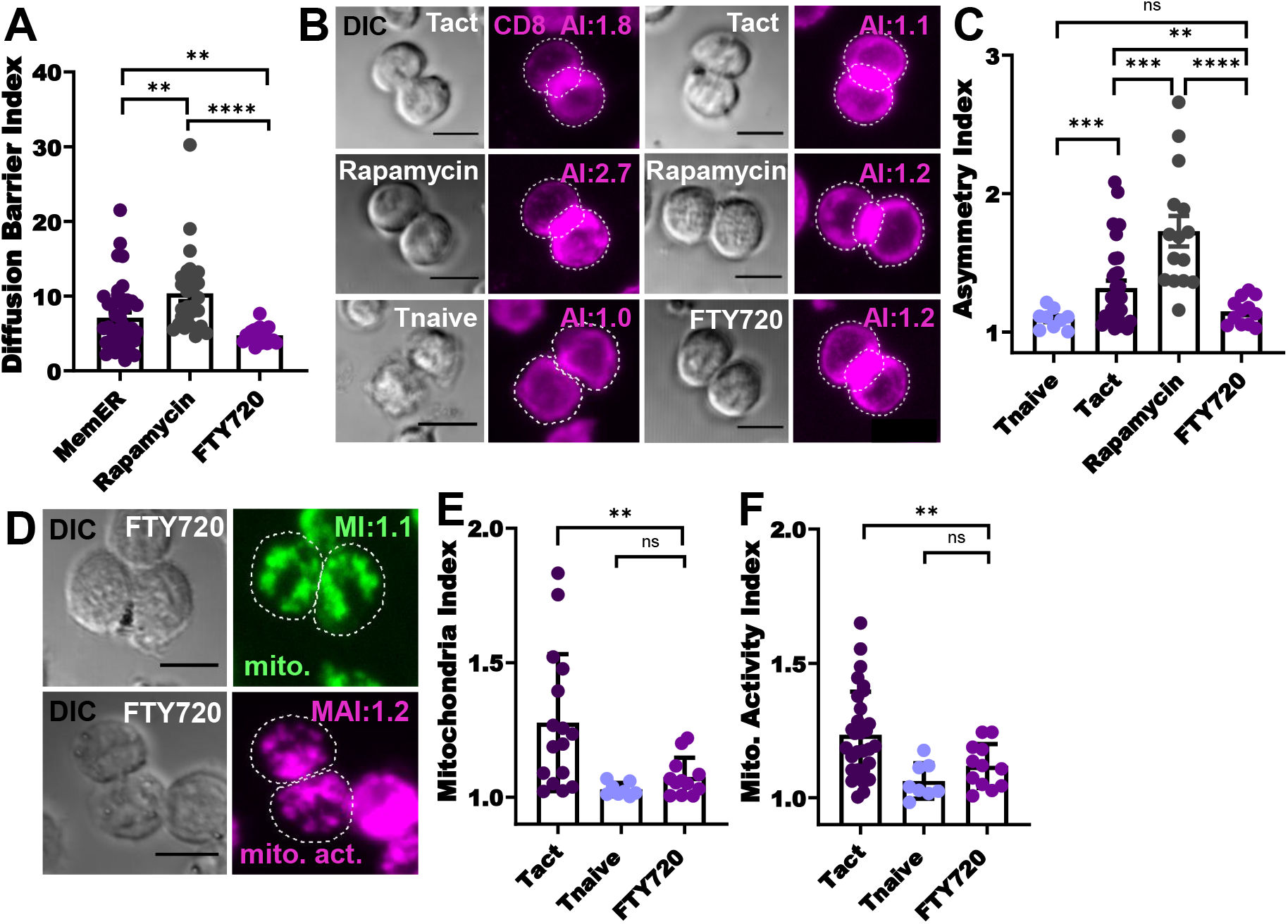
Experimental modulation of the ER diffusion barrier correlates with corresponding changes in phenotypic and metabolic asymmetry of dividing CD8 T cells. FLIP experiments of Sec61-GFP (MemER) transduced TCR activated CD8 T cells performed in presence of rapamycin or FTY720. **(A)** Diffusion barrier strength (BI) in absence of treatment (purple, n=39), in presence of rapamycin treatment (grey, n=27), and in presence of FTY720 (magenta, n=20). **(B)** Representative images of CD8 asymmetry quantification. Naive CD8 T cells induced to undergo homeostatic proliferation were added to the analysis. **(C)** Asymmetry index quantification of activated CD8 T cells (purple, n=30), activated CD8 T cells treated with rapamycin (grey, n=15), activated CD8 T cells treated with FTY720 (magenta, n=12), and homeostatically dividing naïve CD8 T cells (blue, n=9). **(D)** Representative images of MitoTracker Green and Red signals of dividing CD8 T cells treated with FTY720. **(E and F)** Mitochondria index (MI) and mitochondrial activity index (MAI) analysis in sister pairs of dividing CD8 T cells after TCR activation (purple, n=16); after homeostatic cell division (blue, n=8); and after TCR activation in presence of FTY720 (magenta, n=12). Unpaired Welch’s t-test; mean ± SEM. Scale bars, 10 μm. ****P < 0.0001, ***P = 0.0003, **P < 0.05.

### Experimental modulation of ER diffusion barrier strength is related to concomitant alterations in asymmetry of dividing CD8 T cells

To substantiate the notion that the strength of the diffusion barrier is related to the extent of asymmetry established during TCR activated CD8 T cell mitoses, we investigated whether experimental modulation the ER diffusion barrier strength in dividing CD8 T cells has an impact on the asymmetry established between the emerging daughter cells.

Transient *in vitro* exposure of CD8 T cells to either low doses of rapamycin or to FTY720 starting twelve to sixteen hours after activation with agonistic CD3 and CD28 antibodies in presence of ICAM significantly enhanced or reduced the extent of ACD, respectively ^31, 50^. This positive or negative modulation of asymmetry of the first division after activation also had functional consequences, as rapamycin-induced increase of ACD endowed the emerging daughter cells with increased *in vivo* memory potential, while the opposite was the case for the FTY720-treated cells ^31^. We therefore decided to test whether these interventions would also have an impact on the compartmentalization of the ER membrane.

To probe whether rapamycin would enhance diffusion barrier formation in dividing CD8 T cells, we treated naïve CD8 T cells activated by agonistic CD3 and CD28 antibodies in presence of ICAM with low doses of rapamycin starting 16h post-activation. The presence and strength of the barrier were monitored by FLIP microscopy 8 hours later. In addition to evaluating barrier strength, we quantified the distribution of CD8 between the generated T cell sisters by live fluorescence confocal microscopy upon completion of cell division. Rapamycin treatment significantly increased the strength of the diffusion barrier (1.6 fold on average, p<0.05; Fig. 5 A). This was not due to some global slowing down of diffusion in the ER-membrane, as determined by the speed of fluorescence decay in the photobleached compartment, which was not affected by rapamycin (Figure S3). Furthermore, comparison of rapamycin-treated versus untreated T cells indicated that rapamycin treatment stimulated barrier formation in all mitotic CD8 T cells as opposed to untreated cells where a fraction (50%) of cells divided symmetrically. Importantly, rapamycin treatment not only increased diffusion barrier strength, but also promoted asymmetry of CD8 T cell mitoses, demonstrated by the increased asymmetry of CD8 inheritance (Figure 5 B, C).

We noticed that in some TCR-induced CD8 T cell mitoses there was a difference in cell volume of the emerging sister cells, particularly in highly asymmetric mitoses measured by CD8 asymmetry. Combined quantification of cell volume and CD8 fluorescence intensity in sister cells emerging from mitoses induced by TCR activation indeed revealed a strong correlation of CD8 signal intensity and cell volume (Figure S4 A, B). However, CD8 asymmetry was not a mere consequence of different cell volumes, as even more pronounced CD8 asymmetry was observed in mitoses of TCR activated CD8 T cells in presence of rapamycin, where differences in cell volume were abrogated (Figure S4 A, C).

Next, we evaluated whether FTY720 treatment would impair diffusion barrier formation in dividing CD8 T cells. To this end, we treated naïve CD8 T cells activated by agonistic CD3 and CD28 antibodies in presence of ICAM with FTY720 16h post-activation. The strength of the diffusion barrier in the ER membrane was quantified 4 hours later by FLIP microscopy. As with rapamycin, measurement of the speed of fluorescence decay in the photobleached side of the future cleavage plane showed that the FTY270 did not affect the fluidity of the membrane (Figure S3). However, fluorescence decay was much more rapid on the other side of the division plane in the treated compared to the untreated cells. Thus, the diffusion barrier in the ER-membrane was significantly decreased in FTY720-treated cells as compared to untreated cells (Figure 5 A). Notably, FTY20 treatment not only impaired ER compartmentalization, but also led to a strong decrease in CD8 asymmetry (Figure 5 B, C). Furthermore, impairing the formation of the diffusion barrier with FTY720 not only reduced CD8 asymmetry in dividing CD8 T cells, but similarly reduced the asymmetry of mitochondria distribution and activity (Figure 5 D, E), indicating that the establishment of a diffusion barrier in the ER membrane during mitosis is essential to promote metabolic asymmetry in the progeny of dividing CD8 T cells.

As naïve CD8 T cells induced to undergo homeostatic proliferation in absence of TCR triggering failed to induce an ER membrane diffusion barrier (Figure 3 A, G, H), we evaluated whether they would exhibit symmetric inheritance of CD8 and mitochondria. Indeed, inheritance of CD8, mitochondria and mitochondrial activity was symmetric in homeostatically dividing naïve CD8 T cells (Figure 5 B, C, E, F and Figure S3 A-C), comparable to FTY720-treated CD8 T cells activated by agonistic CD3 and CD28 antibodies in presence of ICAM (Figure 5 B, C, E, F).

Taken together, our results indicate that experimental enhancement and impairment of diffusion barrier formation in the ER membrane, respectively, promoted or decreased the division asymmetry of TCR activated CD8 T cells.

## Discussion

This study shows that mitotic TCR activated CD8 T cells are compartmentalized by a lateral diffusion barrier in the endoplasmic reticulum (ER) membrane and that the strength of this barrier positively correlates with the extent of asymmetry imparted to the emerging daughter cells. The strength of the ER diffusion barrier and concomitantly the extent of asymmetric cell division is variable, spanning a spectrum from absence of barrier and asymmetry to strong barrier and asymmetry. The integration of diverse strengths of extrinsic cues are likely responsible for this heterogeneity. Several cell-extrinsic factors were shown to impact on the asymmetry of dividing CD8 T cells: i) the affinity of the agonistic peptide for a given TCR (with high affinity interactions promoting asymmetry ^35^), ii) the presence of integrins promoting the formation and maintenance of the immunological synapse between the antigen presenting cell and the CD8 T cell ^24^, and iii) involvement of TCR triggering in the induction of cell division (induction of cell division by exposure to cytokines (IL-7, IL-15) does not lead to asymmetry (Figure 5). We corroborate that homeostatically dividing CD8 T cells do not establish membrane or metabolic asymmetry and extend this by showing that they also do not compartmentalize their ER-membrane.

Conversely, cell-intrinsic factors that promote or inhibit asymmetry during TCR-induced mitoses of CD8 T cells involve the activity of proteins that regulate cellular polarity such as the atypical protein kinase C (aPKC) isoforms PKCζ and PKC λ/ι), whose inhibition prevented polarization and unequal distribution of IL-2Rα, IFNγR, and T-bet, important factors for effector T cell lineage differentiation. This restulted in impaired memory T cell differentiation but not effector cell differentiation, suggesting that the establishment of ACD is mainly beneficial for securing the differentiation of memory CD8 T cells ^46^. Additional cell-intrinsic pathways involved in the establishment and maintenance of asymmetry are the mTOR pathway (with inhibition of the mTOR pathway promoting ACD ^31^ and memory development ^51^) and the ceramide synthase pathway (with inhibition leading it impaired ACD) ^31^. Here, we corroborate and extend these findings by demonstrating that the same interventions that affect asymmetry of CD8 T cell division also affect the strength of the ER membrane diffusion barrier, strongly suggesting a direct link between these two processes.

Mechanistically, rapamycin was shown to increase ceramide levels both in human and murine cells ^52^ and ceramides are a major component of the ER diffusion barrier in yeast ^53^. Conversely, FTY720 is a ceramide synthase inhibitor, in particular of long chain fatty acid containing ceramides ^54^. The lateral diffusion barrier in the yeast ER is composed of a distinct membrane domain that is thicker (~4 nm) compared to the normal ER membrane thickness (~3 nm), due to the specific enrichment of long fatty acid chain-containing ceramides ^39, 55, 56, 57, 58^. This impedes the lateral diffusion of proteins with transmembrane domain lengths that are common to ER-resident membrane proteins ^57^. Our observation that inhibition of long fatty acid chain ceramide synthesis in dividing CD8 T cells impaired the formation of an ER membrane diffusion barrier lends support to the hypothesis that the ER membrane diffusion barrier in CD8 T cells might also involve long fatty acid chain containing ceramides and it will be interesting to test this hypothesis in further studies.

Functionally, the ability of CD8 T cells to divide asymmetrically was linked to differential fate of the emerging daughter cells. This notion is based on the observation that they differ in phenotype, transcription factor composition, metabolic status, and transcriptional profile ^19, 24, 31, 33, 34, 35^, pre-comitting them to become effector or memory cells ^36^. Also single-cell RNA sequencing of first division daughter CD8 T cells revealed a striking transcriptional dichotomy, reminiscent of pre-effector and pre-memory T cell phenotypes ^36^. In addition, adoptive transfer experiments of first daughter cells emerging after *in vitro* or *in vivo* activation substantiated their differential fate determination. In these experiments, CD8 expression levels of first daughter cells were used to discriminate between cells that (might have) emerged from an ACD. Adoptive transfer of CD8^lo^ first daughters resulted in increased memory potential in comparison to CD8^hi^ daughters ^24, 31, 34^. However, it needs to be stated that this experimental setup using bulk first daugthers with high or low CD8 expression does not allow the unambiguous identification of daughter cells that emerged from an ACD or a symmetric cell division, due to a gaussian distribution of CD8 expression on naïve and activated CD8 T cells. Further evidence for a role of ACD in CD8 T cell fate determination comes from studies where ACD was disrupted or weakened. Cells from ICAM-1 or PKCζ deficient mice show impaired asymmetry upon mitosis, reduced memory T cell responses and biased differentiation towards effector cells ^24, 33, 46^.

Interestingly, CD8 T cells in various differentiation states exhibit different abilities to undergo ACD upon TCR-mediated activation. While naïve and memory CD8 T cells are able to divide asymmetrically ^31, 59^, both cells types being endowed with stemness, i.e. the ability to generate progeny with different fates, end-stage differentiated cells such as effector cells and exhausted cells in the context of chronic viral infection lack the ability to divide asymmetrically^31^. While ACD rates could be increased by transient treatment with rapamycin in both naïve and memory cells upon activation, supporting the generation of offspring endowed with memory potential, this was not the case for end-stage differentiated effector or exhausted cells ^31^. These data support the notion that ACD has a role in assuring diversification of progenies from activated stem-like CD8 T cells. It will be important to address in future studies whether the inability of terminally differentiated cells such as effector and exhausted cells to undergo ACD is linked to an inability to establish an ER membrane diffusion barrier following TCR-mediated activation.

ACD in TCR-activated naïve CD8 T cells was also reported previously to generate asymmetric daughters with respect to their metabolic programmes ^19, 34^. Asymmetric metabolic programmes were installed in ACD daughter cells by unequal mTORC1 kinase activity associated c-Myc, with pre-effector CD8^hi^ T cell daughters exhibiting higher levels of mTORC1 kinase activity, c-Myc expression and glycolysis ^34^ compared to the CD8^lo^ T cell daughters. Interesingly, we observed that in particular very asymmetric CD8 T cell divisions produced first daughter cells with different cell volumes. As the CD8 T cell volume is congruent with anabolic metabolism, this lends further support for diverging metabolic pathways being engaged in first daughter cells from asymmetrically dividing CD8 T cells. In support of this hypothesis, transient inhibition of the mTOR pathway by rapamycin inhibited the generation of first daughter cells with different cell volumes, supporting the notion that differential mTOR activity in first daughter cells is responsible for the difference in cell volume.

In line with these metabolic differences, bulk CD8^hi^ and CD8^lo^ populations of first daughter cells showed unequal inheritance of mitochondrial load ^34^. Unequal mitochondria inheritance via asymmetric cell division was also documented in various different biological systems, such as *Saccharomyces cerevisiae* budding, mammalian oocyte meiosis, and ACD of human mammary stem-like cells ^60^. In this study we also observed unequal partitioning of mitochondria and mitochondrial activity in first daughter cells of dividing TCR-activated CD T cells and we provide evidence that such unequal inheritance is linked to the extent of CD8 asymmetry and the establishment of an ER membrane diffusion barrier. This begs the question how mitochondrial distribution to emerging daughter cells is regulated via the ER diffusion barrier during T cell mitosis. In this context it is noteworthy to mention that mitochondria are fragmented via fission during cell division and this process occurs on ER-mitochondria contact sites ^60^. Therefore, we speculate that mitochondria, in their fragmented form, directly connect to the ER membrane where diffusion can be regulated (to various degrees) by the lateral diffusion barrier that we showed to correlate with asymmetric partitioning of mitochondria during mitosis.

In addition to mitochondria, proteasomes were also shown to be unequally inherited in daughter CD8 T cells during ACD, and prevention of polarization (by interfering with aPKC functions) led to the symmetric inheritance of proteasomes ^46^. Unequal distribution of protesomes was shown to be directly responsible for unilateral degratadion of the master effector cell diffentiation factor T-bet in CD8^lo^ daughter cells ^33^. Interestingly, alike fragmented mitochondria, proteasomes are known to localize to the ER membrane. Therefore, we hypothesize that proteasomal partitioning could also be regulated during T cell division via their association to the ER membrane and be regulated by the establishment of an ER diffusion barrier ^61^. Another mammalian system in which ACD was shown to asymmetrically segregate ubiquitinated proteins, destined for degradation, via the establishment of an ER membrane diffusion barrier preferentially to one of the daughter cells are murine neural stem cell ACD ^42^. Before that, lateral diffusion barrier mediated compartmentalization in asymmetric cell division was characterized in budding yeast ^45^. In yeast, ageing factors such as extrachromosomal DNA or protein aggregates are segregated asymmetrically by the nuclear outer membrane lateral diffusion barrier during yeast budding, permitting rejuvenation of the bud ^38, 39, 45^.

Collectively, our findings establish that depending on the context of their activation, CD8 T cells compartmentalize in a regulated manner their ER through assembly a diffusion barrier in the ER-membrane in the future cleavage plane. The presence of a tight barrier is key for the asymmetric division of these cells upon activation, i.e., the unequal segregation of cell fate and metabolic determinants between daughter cells. Given the importance of such asymmetry in determining the composition and heterogeneity of the progeny, our data suggest therefore that barrier regulation contributes in a decisive manner to establish fate heterogeneity in CD8 T cell responses. The fact that barrier formation and tightness depend on the combination of signals as diverse as TCR and PKCζ activation, TOR signaling and ceramide synthesis puts this process in a central position for integrating context information in order to orchestrate an adapted response. In this respect, the fact that it responds also to metabolic inputs and may contribute to differential metabolic programs in the bifurcating daughter cells opens interesting perspectives for interventions. Further studies are needed to more specifically address the nature of the diffusion barrier in T cells and the mechanism by which it orchestrates the unequal partitioning of specific cellular cargo such as CD8 and mitochondria, but also proteasomes and cell fate-defining transcription factors.

## Materials and Methods

### Animal Experiments

Mice were kept under specific pathogen-free conditions and animal experiments were performed according to the guidelines of the animal experimentation law (SR 455.163; TVV) of the Swiss Federal Government. The protocol was approved by the Cantonal Veterinary Office (ZH115/17 and ZH022/20).

### Constructs

Sec61α-eGFP (MemER-GFP construct) and KDEL-sfGFP (LumER-GFP construct) were a gift from D. Moore, University of Wisconsin-Madison. The genes were cloned into 3rd generation lentivirus vectors under the control of the hPGK (human phosphoglycerate kinase) promoter. pRRL.hPGK.GP91 was a gift from Didier Trono (Addgene plasmid # 30477; http://n2t.net/addgene:30477; RRID:Addgene_30477). Lentiviruses of all constructs were produced as described previously ^42^.

### Naïve CD8 T cell isolation

Naïve CD8 T cells were isolated from young female (6w-8w) B6 mouse lymph nodes and spleens with the EasySep Mouse Naïve CD8 T Cell Isolation Kit (StemCell Technologies) according to the manufacturer’s protocol. T cells were cultured at 37° C / 5% CO_2_ in enriched RPMI medium containing HEPES (25mM; Sigma-Aldrich), 2-Mercaptoethanol (50μM; Sigma-Aldrich), Sodium Pyruvate (1mM; Life Technologies), MEM Non-essential amino acid solution (1X; Sigma-Aldrich), 10% Fetal Bovine Serum (Thermo Fischer Scientific), antibiotics (penicillin-streptomycin; Invitrogen), and the cytokines IL-7 and IL-15 (10ng/ml and 5ng/ml; PeproTech). Such isolated naïve CD8 T cells were cultured for 3d prior transducton with 3rd generation lentiviral vectors on RetroNectin (Takara) coated cell culture plates (according to the manufacturer’s protocol) for 2d without washing. The cells were washed for the first time after an additional at least 5d of culture under the same conditions. At least after 8d of culturing, transduced CD8 T cells were exposed to TCR-mediated activation (anti-CD3, anti-CD28, ICAM), or further exposed to the IL-7 and IL-15 inducing homeostatic cell division, before they were used in FLIP experiments or live confocal fluorescence microscopy.

### Lentiviral transduction

To transduce naive CD8 T cells, culture plates were coated with RetroNectin (20μg/ml) (Takara) at 4o C, overnight, according to the manufacturer’s protocol. CD8 T cells cultured with IL-7, and IL-15 were spun down on RetroNectin coated culture plates in a monolayer with at least 70% confluency. Concentrated lentivirus suspension was gently added onto the CD8 T cell monolayer in a dropwise manner then CD8 T cells were spinoculated at 2000rcf at 30° C for 2h-3h, with acceleration and de-acceleration without a break.

### Activation of CD8 T cells

Cultured and lentivirally transduced naïve CD8 T cells were activated on anti-CD3 (5μM; BioLegend), anti-CD28 (5μM; BioLegend), and ICAM (50μM; R&D Systems) coated 8-chamber live imaging culturing slides (μ-Slide 8 Well; ibidi). Imaging culture slides were incubated with the respective antibodies and ICAM in PBS for 1h at 37° C. CD8 T cells were spun down onto the culture slide surface. The cells were then placed into a 37°C incubator, avoiding shaking the culture. Following 24h of incubation, the culture medium was removed gently from the imaging culture slides without disturbing the monolayer of CD8 T cells. Without letting the cell monolayer drying, T cells were mounted in imaging medium (CyGel mixture: 30% Cygel + 70% enriched imaging RPMI medium with IL-7 (10ng/ml), and IL-15 (5ng/ml)) by gently dropping the solution onto the T cell monolayer, avoiding monolayer detachment from the surface of the chamber slide. Mounted CD8 T cells were then used for FLIP microscopy experiments.

For FLIP experiments of dividing CD8 T cells without TCR activation, naïve CD8 T cells were cultured in enriched imaging RPMI medium (as described previously), without CyGel, with IL-7, IL-15, and IL-2 in 8 chambered live imaging culture slides at relatively high confluency in monolayers.

### Flow cytometry

For flow cytometry-based phenotypic analyses, naïve CD8 T cells were cultured as described previously, washed with cold PBS, and centrifuged. Pellets were suspended and incubated with anti-CD62L (BioLegend), anti-CD127 (BioLegend), anti-CD44 (BioLegend), anti-CD69 (BioLegend), anti-CD25 (BioLegend) antibodies, and LIVE/DEAD Fixable Near-IR stain (Life Technologies), according to the manufacturer’s protocol, and incubated at 4° C for 30 min prior to FACS analysis. Data analysis was performed with FlowJo.

### In vitro live imaging (FLIP experiments)

For *in vitro* FLIP (Fluorescence Loss In Photobleaching) experiments, naïve CD8 T cells were transduced with lentiviruses as described above, and activated on live imaging chamber slides with anti-CD3 (5μM), anti-CD28 (5μM), and ICAM (50μM)). Activated T cells were mounted in a thermo-reversible hydrogel imaging medium as described above. FLIP experiments were performed with a scanning confocal microscope (Zeiss LSM 880) in a compatible cell culture incubator box (EMBL incubator box) at 37° C / 5% CO_2_ conditions. The chambers of live imaging slides were manually scanned for T cells at the mitotic stage and simultaneously monitored with DIC imaging (Differential Interference Contrast microscopy) to detect condensed chromosomes and hence dividing T cells. To perform DIC imaging simultaneously with fluorescence imaging, the DIC slider was inserted underneath of matching imaging objective lens in the correct orientation with each lens having a separate compatible DIC slider. Next, the proper Nomarski prism on the condenser was matched to the DIC slider chosen, and proper Kohler illumination conditions were adjusted on the microscope. To set up for Kohler illumination, we set the focus on the sample with a low power lens using transmitted light (halogen lamp illumination), and the condenser was placed to the lowest position possible. Then, the aperture diagram was opened completely, and the luminous-field diaphragm was closed to make the edges of the sample visible. The condenser was focused to make the edges of the sample as sharp as possible. The image of the luminous-field diaphragm was centered which was followed by proper adjustment of the objective lens, and the luminous-field diaphragm was opened till the edges of the image just outside of the visible region through the eyepieces. Then, the aperture diaphragm was gradually closed to get proper illumination of the view field to complete the Kohler illumination set-up. While scanning, the Wollaston prism was adjusted by the DIC slider to adjust polarization to create proper DIC imaging settings. We followed T cell mitosis through visualizing condensed chromosomal mass via DIC imaging simultaneously, and FLIP experiments were performed for CD8 T cells going through anaphase where condensed chromosomes were just starting to separate. We set the ROI (Region of Interest) for bleaching manually to bleach the specific ER marker, avoiding hitting condensed chromosomes with the laser, since this may interfere with mitosis. A pre-bleach image was recorded for each T cell mitosis, and the ROI was repeatedly photo-bleached throughout the duration of the division. Time-lapse images of FLIP were recorded at a single optical plane after each round a laser hit was completed to monitor fluorescence decay in both future daughter T cells. To confirm that bleaching was performed throughout the T cell, z-stacks (0.5μm step size) were acquired immediately after completion of the FLIP experiment that was performed for each mitosis.

Bleaching experiments were performed with the following settings of fluorescence acquisition: a 63x oil objective (N.A.1.4), acquisitions settings of zoom 2.0, speed 8, 2.5 A.U. (Airy Units), green laser (488nm) at 30% laser power with gain between 700-800, whereas bleaching settings were 70% green laser with 90 iterations using Zen Black 2010 program. Analyses were performed with the ImageJ (Fiji) software package. Fluorescence quantification for future daughters was performed by measuring the area, and RID (Raw Integrated Density) of fluorescence signal for each future daughter of each time-lapse of the FLIP experiment for each ROI and each mitosis. Photobleaching due FLIP timelapse was corrected for each mitosis by control cell photobleaching measurements and background ROI fluorescence intensity correction.

To calculate fluorescence intensity of future daughter cells, the following calculations were performed: ((Bleached (or Unbleached) Compartment RID/Area)-(Bkgrd RID/Area))/((Control cell RID/Area)-(Bkgrd RID/Area)). The pre-bleach image (the first image of FLIP timelapse) was ascribed to be 100% fluorescence intensity, and subsequent frames of time-lapse were calculated related to pre-bleach fluorescence. Collective fluorescence intensity one-phase decay graphs were generated by averaging the fluorescence intensity at a specific frame through FLIP time-lapse of all the cells for both bleached and unbleached future daughter T cell.

For statistical analysis of the one-phase decay curves of fluorescence intensity of the bleached and unbleached daughter cell, the decay curves were used to make an average nonlinear fitted curve. Using Prism software, a one-phase decay was calculated for each FLIP time-lapse of each mitosis with the settings of Plateau=0, Y0=100, and K>0. Statistical analysis data were used to calculate BI (Barrier Index-Barrier Strength), the details of BI calculation were as follows:

- Fit for each future daughter: LN((t-Plateau)/(Y0-Plateau))/(-K)
- Associated standard error of each future daughter: (-SE K*LN(t-Plateau)/(Y0-Plateau))/K^2))
- Barrier Index: (Unbleached daughter Fit/Bleached daughter Fit)
- Barrier Index Standard Error for the Bleached daughter: (Fit Unbleached*total error Bleached / (Bleached Fit ^2))
- Barrier Index Standard Error for the Unbleached daughter: (Total error Unbleached/Fit Bleached)

The calculated barrier index (BI) mean and SE were used to perform a Welch`s t-test with Prism software to determine the Barrier Strength. The calculations and related analyses are based on the related experiments performed in budding yeast (*33*).

FLIP experiments for naïve CD8 T cells were performed with the indicated settings of fluorescence acquisition: a 63x oil objective (N.A.1.4), acquisitions settings of zoom 2.0, speed 8, 2.5 A.U. (Airy Units), green laser (488nm) at 30% laser power with gain between 700-800, whereas bleaching settings were 60% green laser with 90 iterations using Zen Black 2010 program. Analyses were performed with ImageJ (Fiji) software package. Fluorescence quantification for future daughters was performed by measuring the area, and RID (Raw Integrated Density) of fluorescence signal for each future daughter of each timelapse of the FLIP experiment for each ROI and each mitosis. Photobleaching due FLIP time-lapse was corrected for each mitosis by control cells photobleaching measurements and background ROI fluorescence intensity correction.

### In vitro live imaging (scanning confocal microscopy)

To perform and quantify AI (Asymmetry Index) experiments of T cell progeny after completion of the FLIP experiments, anti-CD8 antibodies (Ab) (CD8-APC, BioLegend) were directly pipetted into the hydrogel imaging mixture and incubated for at least 20 min since diffusion of the Ab is slower in the hydrogel in comparison to imaging medium. After completed incubation, z-stacks were acquired with a step size of 0.5μm. Data analysis was performed with ImageJ (Fiji) software package. RID (Raw Integrated Density) of each daughter cell was calculated and normalized with respect to cell area and background signal. The enrichment of the CD8 signal in the contact area in between daughters was excluded since it was not possible to determine the exact quantity of signal that could be assigned to the respective daughters in this area. AI analysis was performed calculating the ratio of each daughter’s CD8 signal.

For experiments that did not involve prior FLIP experiments, AI (Asymmetry Index) experiments and associated analysis in activated T cells were performed as described above.

AI (Asymmetry Index) analysis of homeostatically dividing naïve CD8 T cells was performed without excluding the CD8 signal in the daughters’ cell-cell contact area, since fluorescence signal enrichment in this area was not observed for naïve T cell mitoses under homeostatic proliferation conditions.

### Mitochondrial abundance and activity

Mitochondria Index (MI) and Mitochondria Activity Index (MAI) experiments for both TCR-activated, and naïve CD8 T cells were performed as follows: MitoTracker Green FM (total mitochondria load) (Thermo Fischer Scientific), and MitoTracker Red FM (mitochondrial membrane potential: mitochondria activity) (Thermo Fischer Scientific) were added to CD8 T cells cultured and mounted in hydrogel according to the manufacturer’s protocol. Following at least 30 min of incubation in live imaging chambered slides, CD8 T cell mitoses were manually tracked with DIC imaging simultaneously to fluorescence imaging as described previously. Following mitosis completion, the mitochondrial signals were measured with z-stacks of 0.5μm step size throughout each daughter cell. MI (Mitochondria Index), and MIA (Mitochondria Activity Index) analysis were performed the same way described for AI (Asymmetry Index) analysis of CD8 signal, with the exception of the signal exclusion of cell-cell contact area in between T cell daughters.

### Statistical analyses

Excel (Microsoft) or Prism (Graphpad) were used for all data analyses. Statistical significance was tested with either Welch`s t-test or unpaired t-test with Prism (described in figure legends).

## Acknowledgements

We thank T. Schroeder for scientific discussion, and conceptual input; T. Schwarz, J. Hehl, C. Briand, and G. Csúcs for microscopy technical support; and PPMS for the Scientific Center for Optical and Electron Microscopy (ScopeM) of ETH Zürich. We thank D. Moore for the introduction to FLIP experiments in mammalian cells and for providing FLIP ER reporter constructs. We thank F. Gräbnitz, and N. Barandun for support in flow cytometry.

## Author Contributions

H.E., Y.B. and A.O. designed the experiments; H.E. performed the experiments; Y.B. provided technical assistance; H.E., Y.B. and A.O. analyzed the experiments; H.E., Y.B. and A.O. wrote the manuscript.

## Supplemental Figures

**Figure S1.**
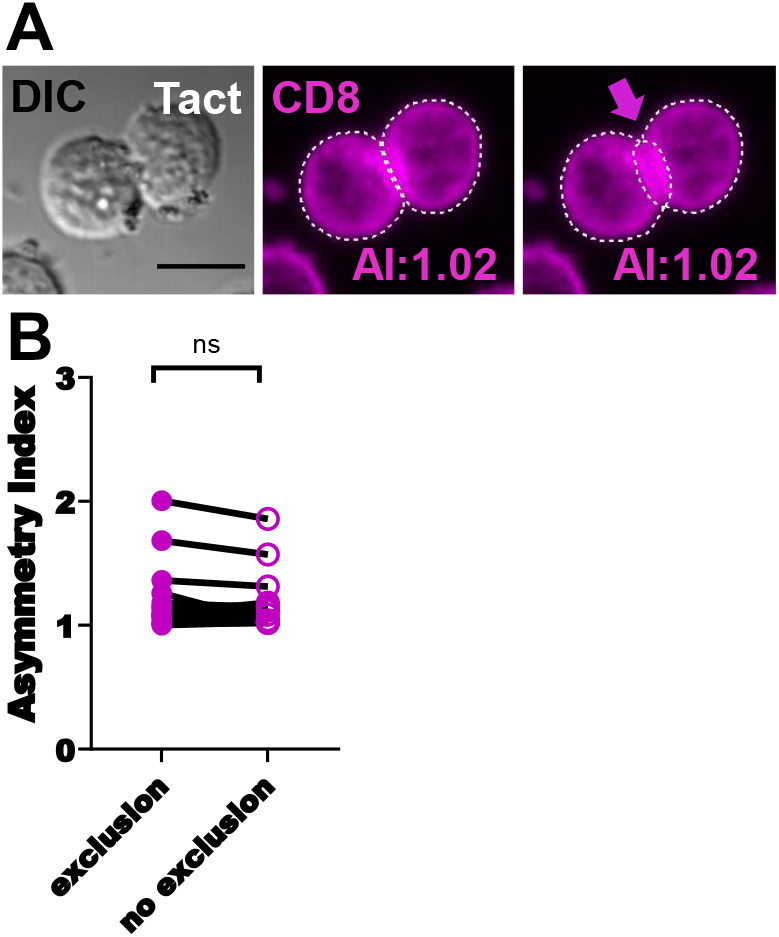
CD8 signal exclusion of cell to cell contact region does not change asymmetry index. **(A)** Representative images of CD8 asymmetry quantification with inclusion or exclusion of CD8 signal at cell to cell contact region. **(B)** Asymmetry index quantification of TCR activated CD8 T cells with CD8 signal of the contact region either excluded or not (pink, n=16). Unpaired Welch’s t-test; mean ± SEM. Scale bars, 10 μm.

**Figure S2.**
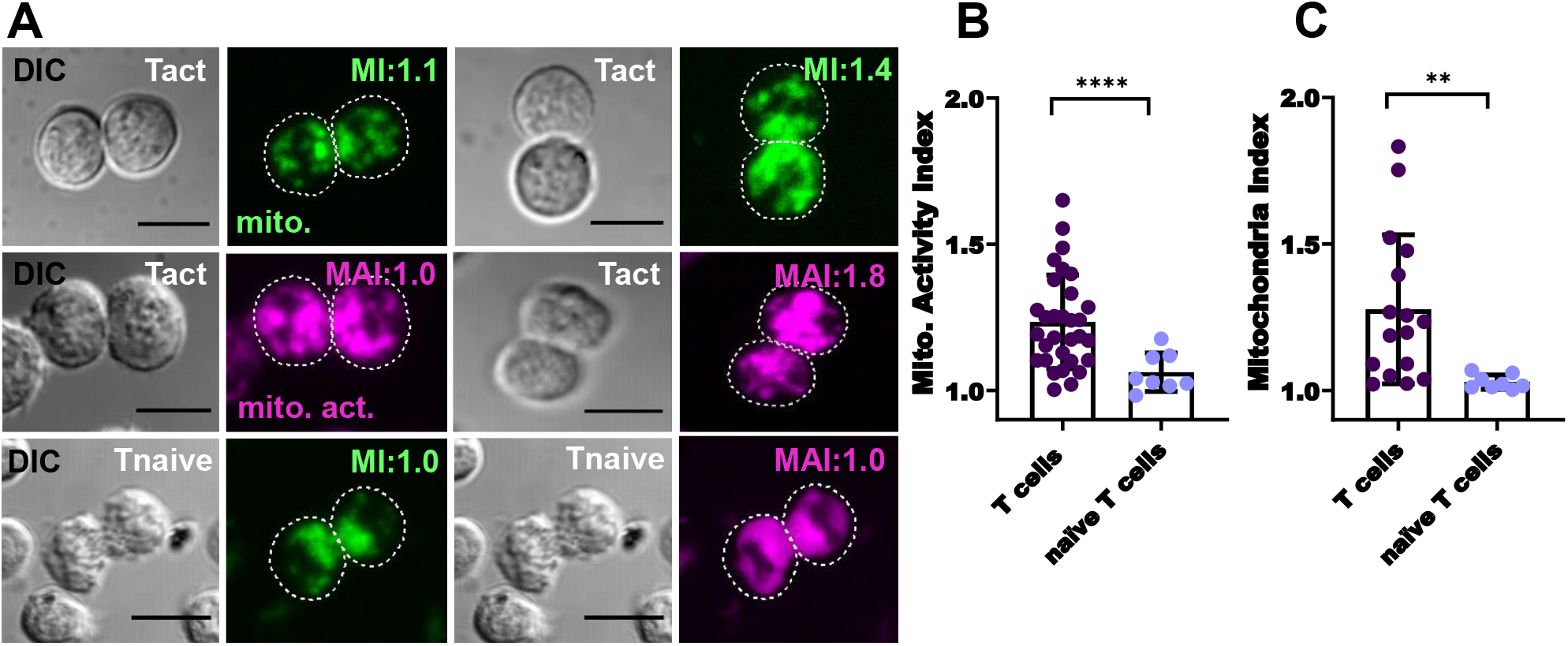
Asymmetric mitochondrial inheritance requires TCR activation. CD8 T cells were either activated by agonistic CD3 and CD28 antibodies in presence of ICAM or cultured in IL-1, IL-15 and IL-2 containing medium to induce homeostatic proliferation. Cells were stained with MitoTracker Green to measure mitochondrial load, and MitoTracker Red to quantify mitochondrial activity 24h post-activation. **(A)** Representative images of live fluorescence confocal microscopy of MitoTracker Green and MitoTracker Red in mitotic CD8 T cells **(B)** Mitochondria activity index (MAI) analysis (ratio of MitoTracker Red signal of sister CD8 T cells after TCR activation (purple, n=31) and after homeostatic division (blue, n=8)). **(C)** Mitochondria index (MI) analysis (ratio of MitoTracker Green signal of sister CD8 T cells after TCR activation (purple, n=16) and after homeostatic division (blue, n=8)). (Unpaired Welch’s t-test; mean ± SEM). Scale bars, 10 μm. ****P < 0.0001, **P < 0.05.

**Figure S3.**
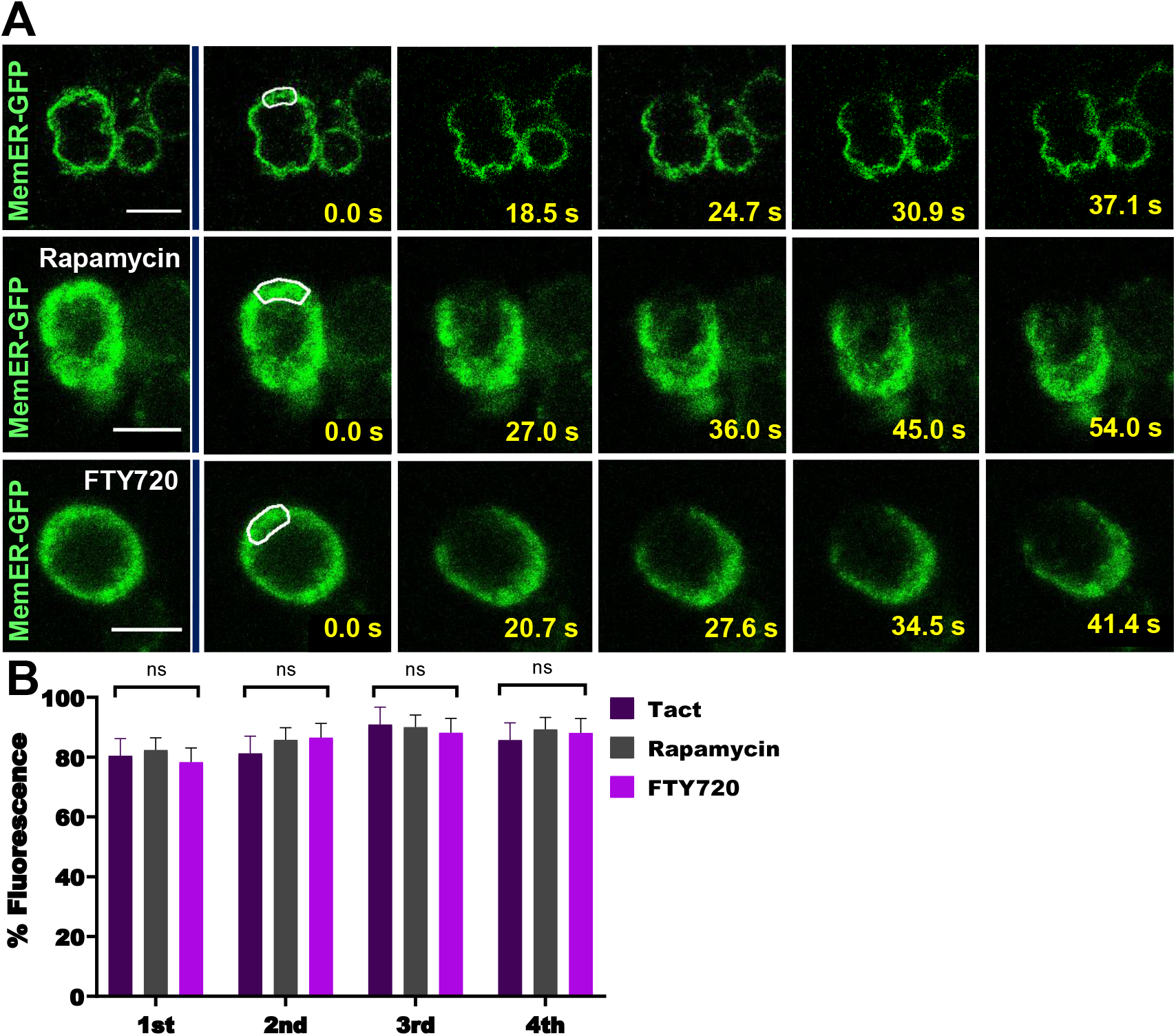
Diffusion of Sec61-GFP does not change with rapamycin and FTY720 treatment. **(A)** FLIP experiments of Sec61-GFP transduced CD8 T cells, activated by agonistic CD3 and CD8 antibodies in presence of ICAM, performed in presence of rapamycin or FTY720 **(B)** Fluorescence decrease quantified in the bleached future daughter cell in four sequential bleaches in the same ROI in untreated cells (purple, n=39), rapamycin treated cells (grey, n=27), and FTY720 treated cell (magenta, n=20). (Multiple t-tests; mean ± SEM). Scale bars, 10 μm.

**Figure S4.**
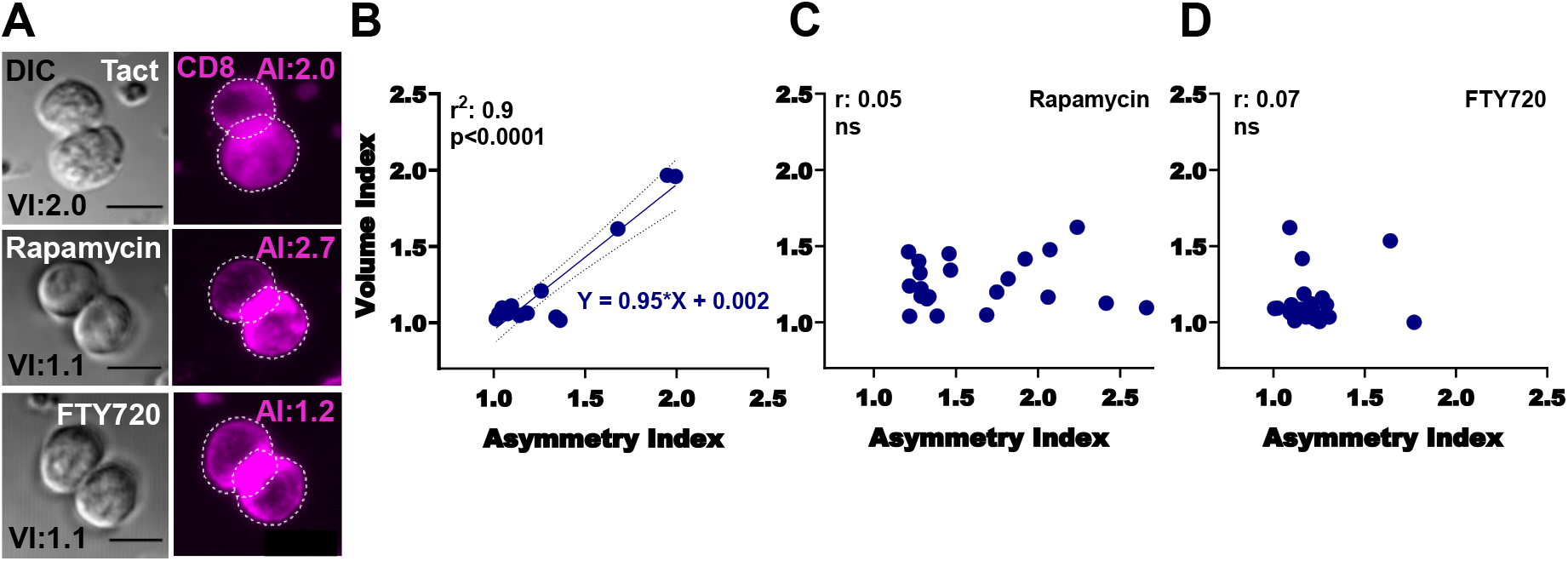
Asymmetric T cell division gives rise to progeny with differential cell volume. **(A)** Representative images of CD8 asymmetry quantification and cell volume quantification of sister CD8 T cell pairs. Asymmetry and cell volume index (VI) quantification of activated CD8 T cells, activated CD8 T cells treated with rapamycin, and activated CD8 T cells treated with FTY720. Linear regression of the asymmetry index and associated cell volume index (the ratio of cell volumes of the sister T cell pairs) of a particular mitosis’ progeny in absence of treatment (**B**, n=15) in presence of Rapamycin treatment (**C**, n=21), or in presence of FTY720 treatment (**D**, n=21).

